# Ultra-high throughput *in vitro* translation assay for identification of *Trypanosoma cruzi-specific* protein synthesis inhibitors

**DOI:** 10.1101/2025.02.02.636133

**Authors:** Camila Colombari Mantovani, Carolina Borsoi Moraes, Guilherme Rodrigo Reis dos Santos, Bertal Huseyin Aktas, Sergio Schenkman

## Abstract

Chagas disease, a global health threat exacerbated by the migration of infected individuals, raises significant concern due to its restricted treatment options, resulting in around 12,000 fatalities annually. The disease is caused by *Trypanosoma cruzi,* a protozoan parasite. *T. cruzi* possesses a divergent translation apparatus, characterized by enlarged ribosomes and a unique mRNA cap appended to the 5’ end of a 39-nucleotide spliced leader inserted upstream of all its mRNAs. Given that many anti-microbial agents are protein synthesis inhibitors and the divergence of *T. cruzi* protein synthetic machinery compared to its mammalian host, it is likely that *T. cruzi-specific* protein synthesis inhibitors can be employed for the treatment of Chagas disease. Such inhibitors can be identified by screening chemical libraries in an *in vitro* translation assay. However, lysates of *T. cruzi* cannot reinitiate translation of exogenous mRNAs. Here, we demonstrate that *T. cruzi* extracts derived from a strain deficient for hemin accumulation efficiently translate a capped and polyadenylated reporter mRNA. We optimized this assay’s conditions and adapted it to a 384-well format. In a proof of principle pilot study, we identified quinazoline compounds that inhibit translation of reporter mRNAs *by T. cruzi* extracts. These compounds inhibited protein synthesis in live wild-type parasites. We further miniaturized this assay to a 5 µL volume, demonstrating its suitability for ultra-high throughput screening in a 1536-well format with appropriate automation. This first-of-its-kind *T. cruzi ultra-high-throughput in vitro* translation assay will enable screening of very large chemical libraries while hit compounds identified in the pilot study can be developed into lead compounds by synthesis and/or assembly of focused libraries to develop therapeutics for Chagas disease.

## Introduction

Chagas disease, originally endemic to Latin America, has been globally disseminated due to the emigration of infected individuals and the expanding range of insect hosts, resulting in a significant public health concern around the world. Chagas disease accounts for >12,000 fatalities annually, with around 75 million individuals at risk of infection (DNDI, 2021; World Health, 2021). The disease is caused by the protozoan *Trypanosoma cruzi*, which is transmitted by blood-sucking insect vectors, through blood transfusion and organ transplants from infected patients, or by congenital contact (Coura, 2015; Shikanai Yasuda, 2022; Zingales & Bartholomeu, 2022). *T. cruzi* multiplies in the midgut of the insect host as epimastigotes and transforms into non-proliferating but infective metacyclic trypomastigotes, which are released with the feces or urine during the blood meal. They reach mucosal membranes of mammalian hosts and enter cells, forming a parasitophorous vacuole, then escaping to the cell cytosol where they proliferate in amastigote form. Amastigotes differentiate into trypomastigotes that are released after lysis of the infected cells, reaching the extracellular environment to disperse through different tissues and cell types. Upon human infection, the parasite causes various symptoms, including headache, body aches, and nausea. Infection typically self-resolves due to a robust immune response. However, a persistent infection remains for decades, and in some cases heart or digestive tissues start to present alterations, most likely due to imbalances in the host’s immune system leading to cardiac, gastrointestinal, and neurological complications, with potential death.

Two drugs are available for the treatment of Chagas disease: Benznidazole and nifurtimox (Coura & Borges-Pereira, 2011; Hasslocher-Moreno et al., 2021; Kratz, 2019; Kratz et al., 2022; Urbina, 2015). They are prodrugs, requiring metabolization by the parasite to become active, and are primarily useful during the acute phase of the disease (Maguire et al., 2021; Wilkinson et al., 2008). The treatment is complicated due to drug cytotoxicity and induction of several side effects, including nausea, sensory neuropathy, fever, muscular and joint discomfort, and weight loss, among others, with the worst side effects among the elderly, causing them to interrupt the treatment (Maguire et al., 2021). During the chronic symptomatic phase, drugs are prescribed according to the patient’s medical records and the disease evolution, but chances of cure are low, and some strains can develop resistance (Campos et al., 2017; Diaz-Viraque et al., 2018; Murta et al., 2008; Porta et al., 2023).

The mechanism of action of numerous antimicrobial medicines employed in the treatment of infectious disorders is the inhibition of protein synthesis (Coyle & Society of Infectious Diseases, 2003; Hilal-Dandan & Brunton, 2016). Despite *T. cruzi* being an eukaryote, its translation machinery shows a significant divergence from mammalian protein synthesis. *T. cruzi* genomes are organized in polycistronic clusters that are transcribed as one long pre-mRNA that undergoes trans-splicing to add a peculiar 39-nucleotide sequence named spliced leader (SL) to its 5’ end. In addition to the 7mGTPpp cap present in mammalian mRNAs, in *T. cruzi* mRNAs, the next 4 nucleotides are also methylated on their base, generating a Cap4 structure. (Liang et al., 2003; Sather & Agabian, 1985; Zwierzynski & Buck, 1991). *T. cruzi* translation initiation factors and ribosomal RNA are also different from their mammalian counterparts (Gao et al., 2005; Liu et al., 2016; Zinoviev & Shapira, 2012). These differences are large enough to be exploited for the development of new drugs for the treatment of Chagas disease, ultimately resulting in its cure.

Identifying inhibitors of *T. cruzi* protein synthesis by screening large chemical libraries in an *in vitro* translation assay would be a straightforward affair. However, we have previously found, and others reported, that lysates of *T. cruzi* cannot synthesize large proteins (Grossi de Sa et al., 1984), and our observations indicate it coul reinitiate translation of exogenous mRNAs. Based on the accumulation of hemin in *T. cruzi* vacuoles, we hypothesized that excess hemin or molecules derived from or modified by hemin during lysate preparation inhibit re-initiation by *T. cruzi* extract. Herein we demonstrate that extracts derived from a *T. cruzi* strain deficient for hemin accumulation efficiently translate capped and polyadenylated reporter mRNAs. We optimized this assay’s conditions and adapted it to a 384-well format. In a proof of principle, we identified quinazoline compounds that inhibit translation of reporter mRNAs by *T. cruzi* extracts. These compounds inhibit protein synthesis in live wild-type *T. cruzi*. Finally, we miniaturized this assay to a 5 µl volume, demonstrating its suitability for ultra-high throughput screening in a 1536-well format with appropriate automation.

## Materials and Methods

### Compounds

The compounds used in the assays were obtained from a Broad Institute library previously tested for inhibition of *Trypanosoma cruzi* proliferation (PubChem AID 2294), from the Drugs for Neglected Diseases initiative, Latin America (DNDi), purchased from Molport (Lithuania) or from the author’s in-house collections at Harvard Medical School (Boston, MA). All compounds were resuspended in 100% DMSO to a stock concentration of 10 mM, and aliquots were stored at −80°C to avoid freeze-thaw cycles. The list of compounds is presented in the supplementary table S1. The tests were performed at a final concentration of 40 μM, unless otherwise indicated. Analogs with >80% similarity to the most promising compounds were purchased from Molport (see table S1).

### Parasite extracts

*T. cruzi* epimastigote form of wild type (WT, DM28c strain) and genetically modified parasites of the DM28c strain (TCH) were maintained at 28°C in LIT medium (75 mM NaCl, KCl 5.4 mM, Na_2_HPO_4_ 62 mM, glucose 0.2%, tryptose 0.5%; liver infusion 0.5%, hemin 10 μg/mL), containing 10% fetal bovine serum (FBS), 59 μg/mL penicillin, and 133 μg/mL streptomycin, adjusted for pH 7.2 (Camargo, 1964). The procyclic form of *Trypanosoma brucei* 29-13 was maintained in SDM-79 media as described in (Machado et al., 2018). The TCH strain contained the genes of T7 RNA polymerase and Cas9 protein of *Streptococcus pyogenes* inserted in the pTREX vector (Beneke et al., 2017). It corresponded to a parasite clone selected in the presence of 250 μg/mL of hygromycin aiming to replace the Vps23 gene of the endosomal sorting complexes required for transport (ESCRT) machinery. The clone was produced by transfection in an Amaxa nucleofactor (program X14) with two sgRNA DNA templates and a donor DNA that contained the hygromycin gene flanked by 5’ and 3’ corresponding to the Vps23 gene (see description in the supplementary section).

The extracts for *in vitro* translation were prepared based on the protocol developed by Kovtun and collaborators, described for *Leishmania tarentolae* (Kovtun et al., 2011). Briefly, *T. cruzi* cultures in the log growth phase (2 to 3 x 10^7^/mL) in LIT-10% FBS medium was collected by centrifugation at 2,500 x *g* for 20 minutes at 4°C, then resuspended and washed twice by the same centrifugation conditions in sucrose elution buffer (SEB, composed of 45 mM HEPES-KOH pH 7.6, 250 mM sucrose, 100 mM potassium acetate, and 3 mM magnesium acetate, all from Sigma-Aldrich) at 4°C. The final pellet was then resuspended in SEB to a concentration of 1 x 10^10^ parasites/mL. These parasite suspensions were lysed by nitrogen cavitation under pressure of 70 bar of nitrogen for 120 minutes at 4°C. The resulting lysates were centrifuged at 10,000 x g at 4°C for 20 minutes, and the recovered supernatant was centrifuged at 30,000 x *g* at 4°C for another 20 minutes. Each 3 mL of the supernatant was applied in a 1 x 8 cm column of Sephadex G-25 (Cytiva) pre-equilibrated with the same solution without sucrose (EB). The first 3 mL were discarded, and the next 4 mL eluted with EB were retained. The extracts were quick-frozen in dry-ice ethanol in 500 μL aliquots and stored at −80°C. An aliquot was used for determining protein concentration using the Pierce BCA Protein Assay Kit (Thermo Fisher Scientific, USA), following the manufacturer’s instructions.

### RNA preparation

A synthetic DNA sequence comprising the SP6 promoter, 39 base pairs corresponding to the *T. cruzi* spliced leader sequence, followed by 34 randomly assigned base pairs, and the Renilla luciferase open reading frame, along with 306 base pairs of the 3’ UTR of *T. cruzi* β-tubulin, was inserted into the KpnI and NotI sites of the pBSK+ plasmid (Stratagene). The pBSK plasmid was amplified in *Escherichia coli* DH5α and purified using the *illustra™ plasmidPrep Midi Flow Kit* (GE Healthcare, USA). For RNA synthesis, plasmids were linearized with *NotI* (Thermo Fisher Scientific) and repurified with the *PureLink Quick PCR Purification Kit* (Invitrogen).The transcription reactions were carried out with HiScribe^®^ SP6 RNA Synthesis Kit (New England Biolabs, USA) in a final volume of 20 μL with 1 μg of template DNA following the manufacturer’s instructions. The generated RNAs were recovered and quantified using the NanoDrop 2000 spectrophotometer (Thermo Fisher Scientific, USA). 10 µg of RNA was capped using the *Vaccinia Capping System* (New England Biolabs, USA), followed by the addition of a polyadenine tail at the 3’ end by the *E. coli Poly(A) Polymerase* (New England Biolabs), according to the manufacturer’s recommendations. The modified RNA was analyzed by Tape Station (Agilent Technologies) and stored in aliquots containing 80% ethanol at −80°C.

### Translation assay

Lysate aliquots (500 μL) were thawed and supplemented with 200 µL of the *feeding* solution according to (Kovtun et al., 2011) [6 mM ATP, 22.5 mM Mg(OAc)_₂_, 200 mM phosphocreatine, 200 U/mL creatine phosphokinase, 57 µL amino acids mix solution, 100 mM HEPES-KOH pH 7.6, 5% (v/v) Polyethylene glycol 3.350 from Sigma-Aldrich, 0.68 mM GTP, 1.25 mM spermidine, 10 mM dithiothreitol (DTT) from Thermo-Fisher Scientific, and 5x protease Inhibitor Cocktail (*Complete*™ *EDTA-free*) from Roche, Switzerland]. The reactions were performed in triplicate in a 384-well white plate, with a final volume of 20 μL containing 14 μL of the supplemented lysate and 2 μL of modified RNA in each well. As a negative control, reactions were conducted without RNA or 10 μM of hygromycin. To test protein synthesis inhibitors, 14 μL of supplemented lysate was mixed with 4 μL of each compound, diluted in assay buffer (final concentration of 40 μM), centrifuged, and pre-incubated at room temperature for 15 minutes. After adding the RNA, the reaction was incubated for 2 hours at 28°C.

The Renilla Luciferase Assay System (Promega, USA) was employed to measure the luciferase expression, according to the manufacturer’s instructions. The luminescence detection, measured in relative light units (RLU), was made immediately after the translation reaction upon the addition of 50 μL of reagent per well in the SpectraMax M3 Microplate Reader (Molecular Devices LLC, USA) with a 500 ms integration time. To rule out the possibility that the compounds are direct inhibitors of Renilla luciferase, the initial hit compounds were added to the mixture after the translation reaction, before the addition of the Renilla reagent.

### Assay miniaturization to low-volume 384-well format

We adapted and optimized the above assay to low reaction volumes (8 µL and 5 µl)) to test multiple reagents with the best possible sensitivity. Extracts supplemented with the feeding solution remained the same. In each well were placed 3 μL of the supplemented lysate (2.6 μg protein/μL) and 2 μL of each compound (40 μM final) pre-incubated for 15 minutes at room temperature. Then 3 μL of RNA (9 ng total) was added, and the reactions were incubated for 2 hours at 28°C. After incubation, 8 μL of Renilla luciferase reagent was added, followed by light detection. For the miniaturized assays, we used 2 μL of the supplemented 1 μL of the supplemented lysate.

### Puromycylation assay and Western Blotting

*T. cruzi* epimastigotes of DM28c (WT) maintained at 28°C in LIT medium containing 10% FBS at 1 x 10^7^ parasites/mL were centrifuged at 2,000 x *g* for 5 minutes, washed, and resuspended in LIT-10% FBS to the 1 x 10^7^ parasites/mL concentration. The parasites were then pre-incubated at 28°C for 1 hour after adding 1 µL of each compound pre-diluted in DMSO for each 1 mL parasite suspension to obtain a concentration of 40 µM. With this concentration of DMSO, we observed no changes in *T. cruzi* motility and viability measured using a Guava Muse cell analyzer. The controls included incubation with DMSO and 5 μM cycloheximide. Thereafter, 10 μg/mL puromycin (Thermo Fisher Scientific) was added to all tubes, which were maintained at 28°C for 15 minutes, followed by centrifugation at 2,000 x *g* for 5 minutes, washing with PBS, and lysis with 50 µL of RIPA buffer (50 mM Tris, 150 mM NaCl, 0.5% sodium deoxycholate, 1% NP-40, 0.1% SDS pH 8.0). 10 μL were used to measure the protein content using the Pierce BCA Protein Assay Kit (Thermo Fisher Scientific, USA). The remaining solution was mixed with SDS-PAGE sample buffer (4% SDS, 20% glycerol, 10% 2-mercaptoethanol, 0.004% bromophenol blue, 120 mM Tris HCl, pH 6.8), heated at 95°C for 5 minutes, and 20 µg protein was loaded into a 10% SDS-PAGE. After electrophoresis, the proteins were transferred to nitrocellulose membranes using a semidry Bio-Rad transblot at 20 volts for 40 minutes. After staining in 0.5% Ponceau S in 1% v/v acetic acid, the membranes were soaked in 5% BSA in 10 mM Tris-HCl pH 7.4 (TBS). After blocking, the membrane was washed three times with TBS with 0.05% Tween-20 (TBS-Tween) and incubated with anti-Puromycin antibody (Mouse ab, PMY-2A4-s, Developmental Studies Hybridoma Bank) diluted 1 to 100 in 5% BSA-TBS containing 0.02% NaN_3_ overnight at 4°C. Following 3 x 10-minute washes in TBS-T, bound antibodies were detected by incubation with IRDye680 goat anti-mouse IgG diluted 1 to 10,000 in TBS-T, washed again, and visualized using Odyssey Fc Imager (LI-COR Biosciences - USA). As a loading control, we used a rabbit anti-aldolase antibody (Barbosa Leite et al., 2020), diluted 1 to 2000.

### Statistical analysis

The statistical analysis was made using GraphPad Prism Software (10.4.1).

## Results

### Establishment of optimal conditions for in vitro translation by T. cruzi extracts

We initially tested the capacity of wild-type extracts made from *T. cruzi* epimastigote cultures prepared as described for *L. tarentolae* to translate an mRNA containing the spliced leader and 5’UTR sequence fused to the Renilla luciferase open reading frame and a 3’ UTR of the *T. cruzi* β-tubulin, without success. It was intriguing that by using *Trypanosoma brucei* procyclic extracts made in parallel, we could generate active luciferase (data not shown). A characteristic of *T. cruzi* epimastigote cultures is the accumulation of hemin within the parasite, as a growth requirement, which results in brownish fractions during the extract preparations. Recently we obtained a mutant parasite (TCH), which accumulated much less hemin due to its incapacity to adequately perform endocytosis. Considering that the presence of hemin might interfere with reinitiation in the extracts, we decided to test the capacity of an extract prepared from this mutant to perform translation. In fact, the resulting parasite extract translated capped and polyadenylated reporter mRNA significantly more efficiently compared with the extract of wild-type parasites (Figure 1A). Both extracts contained similar amounts of protein (Figure 1B), but the wild-type extracts presented a yellow color, probably resulting from the presence of high hemin concentration.

**Figure 1.**
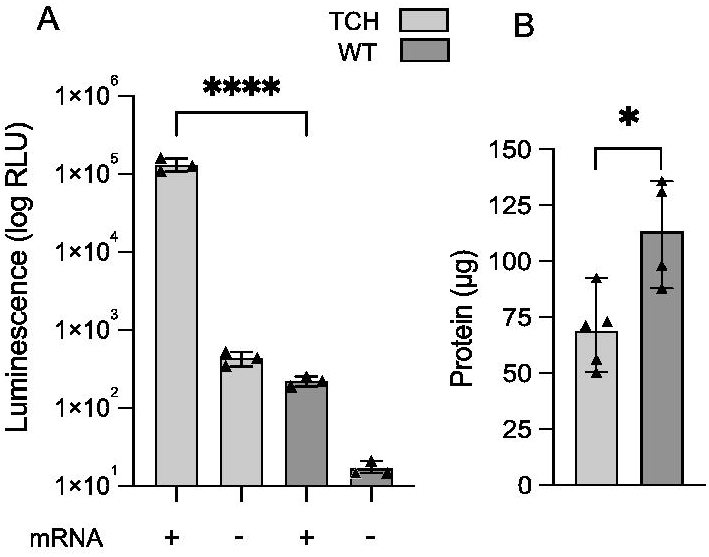
A cell extract from a modified *Trypanosoma cruzi* line is able to promote *in vitro* translation of Renilla. (**A**) Renilla luciferase activity obtained by 3 hours at 28°C incubation of a supplemented extract from a modified (TCH) and original DM28c strain (WT) in the absence (-) or presence of 3 ng/μL capped and polyadenylated Renilla luciferase containing the SL sequence (+), followed by the addition of Renilla luciferase reagent (Promega) and luminescence detection. (**B**) Total protein content used in TCH and WT extracts, quantified by BCA.

To establish ideal conditions for the assay, we then tested different RNA concentrations, incubation times, TCH extract concentrations, and mRNA modifications. The translation of unmodified *in vitro* transcribed RNA was almost the same as background. Capping the mRNA or capping and polyadenylation, dramatically increased the translation (Figure 2A), demonstrating the importance of those modifications to the parasite’s translation machinery. The luminescence signal was directly proportional to the concentration of fully modified RNA up to 3 ng/μL (Figure 2B) with a linear signal up to 2 h reaction time (Figure 2C). Translation was also linear by up to 3 µg/mL of the supplemented parasite extract (Figure 2D). A decline in translation was observed at higher protein concentrations. These optimal conditions were then used in the subsequent experiments.

**Figure 2.**
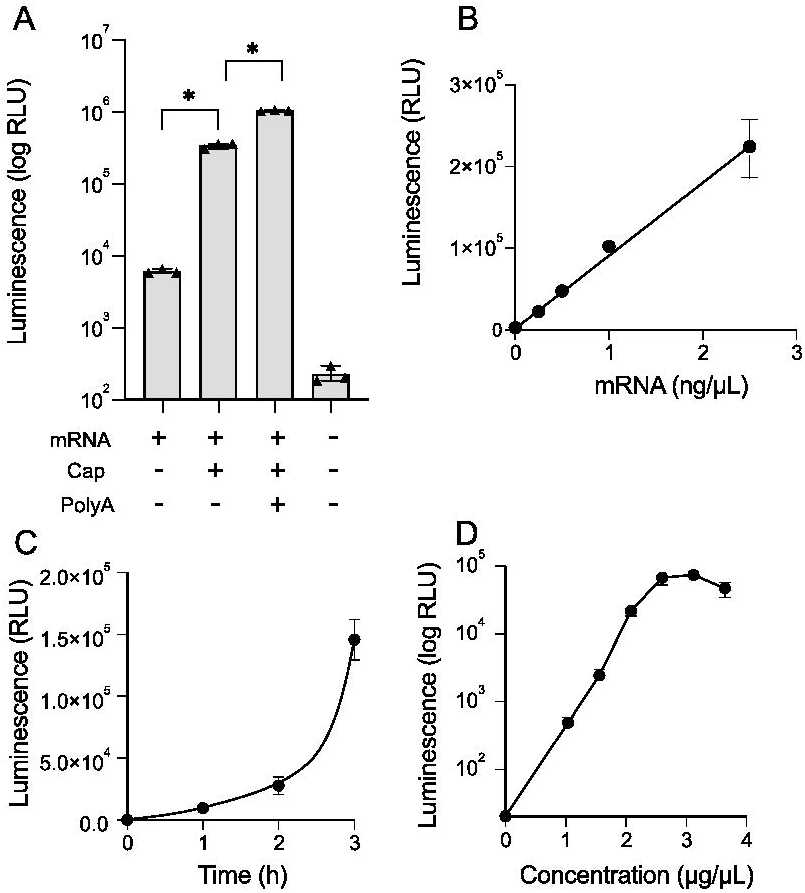
Effect of mRNA modifications, concentration, time of incubation, and extract concentration on Renilla luciferase translation. (A) Renilla luciferase activity using 20 ng of SP6-transcribed RNA (mRNA), before or after capping (Cap) or polyadenylation (PolyA) modifications after 3 hours with supplemented TCH extract, followed by relative luminescence units (RLU) detection. (B) RLU of translation reactions with different concentrations of capped and polyadenylated mRNA incubated with supplemented TCH extract. (C) RLU of translation reactions of 20 ng of capped and polyadenylated mRNA incubated for the indicated times with supplemented TCH extract. (D) RLU produced after translation using different extract concentrations with 20 ng of capped and polyadenylated mRNA for 2 hours. The results are means and standards (SEM) of means of triplicate experiments, each made in triplicate samples. Asterisks indicate significant differences (p < 0.05) using a two-way ANOVA test.

We next validated this assay by demonstrating that known general protein synthesis inhibitors reduced translation when tested at 10 μM (Figure 3A). Paromomycin, a protein synthesis inhibitor used to treat visceral leishmaniasis (Coser et al., 2020; Pokharel et al., 2021), did not inhibit translation in *T. cruzi* extract. Because TCH parasites were partially resistant to hygromycin B, we evaluated its effects in TCH and WT extracts. Hygromycin similarly suppressed translation by the TCH extract (Figure 3B), though slightly less efficiently than WT, indicating that it can be used as a control to screen anti-*T. cruzi* compounds.

**Figure 3.**
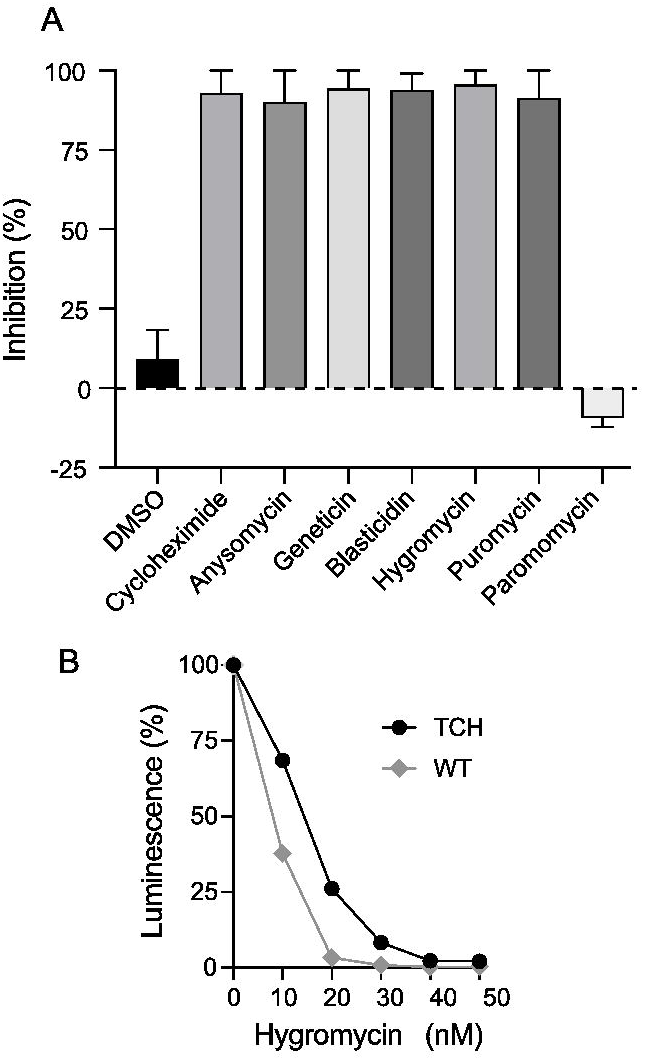
Translation of Renilla luciferase is inhibited by known protein synthesis inhibitors. (**A**) Capped and polyadenylated Renilla mRNA (20 ng) was incubated with the supplemented TCH extract in the presence of 10 μM of the indicated compounds for 2 hours, followed by luminescence detection. The values show the percentage of inhibition relative to the RLU produced in samples containing DMSO. (**B**) Hygromycin B dose-response curve using WT and TCH extracts. The same concentration of TCH and WT-supplemented extracts was incubated with mRNA (20 ng) and different hygromycin B concentrations for 2 hours, followed by luminescence detection. The values indicate the luminescence produced for each extract relative to the values in the absence of hygromycin B. All values are means and SEM of triplicate experiments.

### Proof-of-concept pilot screen

We reasoned that protein synthesis inhibitors should be enriched among the compounds that inhibit protozoa proliferation. We therefore assembled a small library of such agents rather than test a larger random library as a proof of concept demonstration. We tested this 128-member library (Table S1) at a final concentration of 40 μM (see materials and methods). Of these, eleven compounds reproducibly reduced Renilla luciferase signal in the *in vitro* translation assay, ranging from 20% (C3, C39, C124, and C125) to nearly 100% (C108) (Figure 4A). To determine whether the compounds were bona fide inhibitors of translation rather than enzymatic activity of Renilla luciferase, we incubated the extract with these 11 compounds either before or after the two-hour translation reaction and measured the Renilla luciferase activity. The bona fide inhibitors of translation will inhibit luciferase activity only if added to the translation reaction as described in the material and methods section, while inhibitors of enzymatic activity of the Renilla luciferase would inhibit luciferase signal regardless of whether they are added to the reaction before the start of the reaction or after the reaction is terminated. Compounds C3, C39, and C97 appear to inhibit translation (Figure 4B). In contrast, C48, C77, and C108 appeared to inhibit the enzymatic activity of the Renilla luciferase. Compounds C25 and C43 appeared to have a significantly higher effect on the enzyme activity if added before the start of translation but also appeared to inhibit the enzymatic activity of Renilla luciferase when added after the reaction.

**Figure 4.**
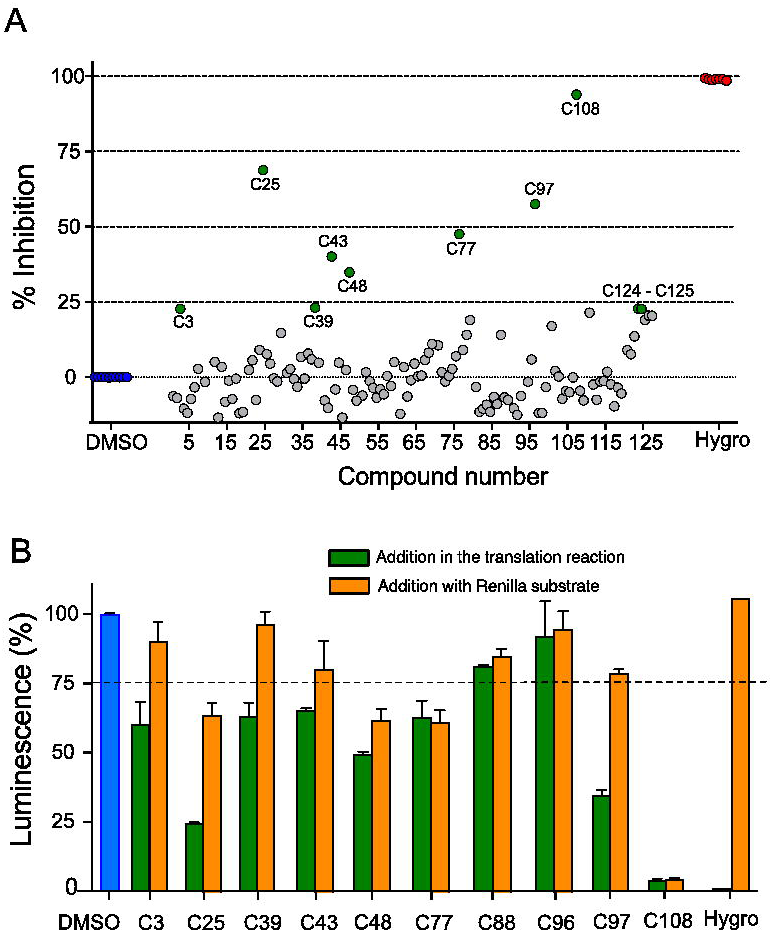
*In vitro* translation inhibition by compounds known to affect parasites. (**A**) Capped and polyadenylated Renilla mRNA (20 ng) was incubated with the supplemented TCH extract in the presence of 40 μM of the compounds named on Supplementary Table S1 for 2 hours, followed by luminescence detection. The values show the triplicate means of the percentage of inhibition of three independent experiments relative to the RLU produced in samples containing DMSO. The significant inhibition (p < 0.05 using two-way ANOVA and Tucker test) is shown in green symbols. The inhibition by hygromycin at 10 μM is shown in red symbols. (**B**) Luminescence values obtained with the same reaction as in (A) with the compounds added before the translation reaction (green bars) or after the translation reactions relative to the values of DMSO (blue bar). The values are means and SEM triplicate experiments. Compounds that were added after translation and which produced less than 75% luminescence were considered Renilla inhibitors.

### Miniaturization of assays to low volume and dose response studies of hits and their analogs

To demonstrate the utility of the miniaturized assay for testing, we utilized it for testing the analogs of the *T. cruzi* protein synthesis inhibitors identified as described above. Due to the structural similarity between quinazolines C39 and C97, we choose to assess analogs of these compounds and compound 25 using the 8 μL reaction volume (Figure 5A). The quinazoline analogs 39A, 39B, 39D, 39G, 39H, 39I, and 39K reduced translation by > 50%, whereas 39C, 39F, and 39L exhibited about 50% inhibition, and 39M showed less than 40% inhibition (Figure 5B). We selected three quinazolines (39, 39D, and 39L) to conduct a dose-response curve, utilizing doses ranging from 2.5 μM to 40 μM (Figure 5C). Compound 39 exhibits a considerable inhibition effect at 30 μM, but the other compound achieves the same rate at 40 μM. In the initial tests, the quinazoline compound 97 was more than 2X more active than 39. Unfortunately, we could not test compound 97 in these dose-response studies because this compound is no longer commercially available. These data indicate that structural modifications influence the efficacy of the quinazolines in inhibiting *T. cruzi* protein synthesis.

**Figure 5.**
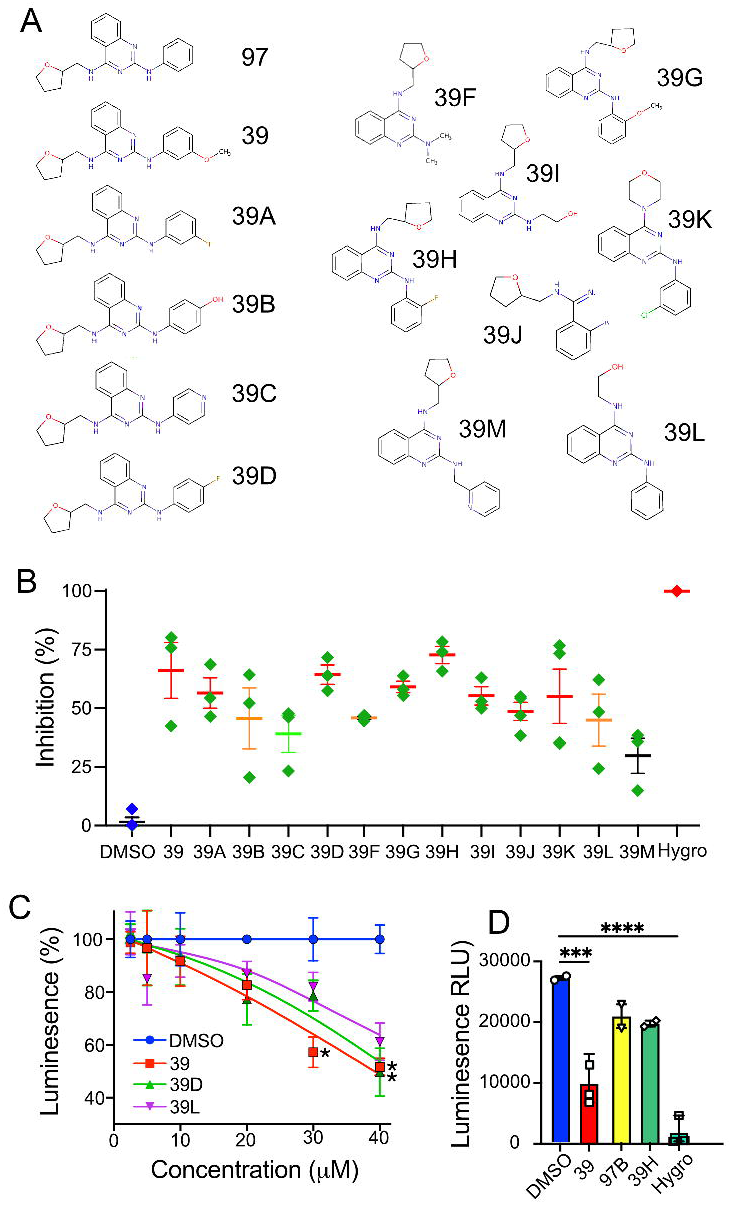
Effect of compound analogs on the *in vitro* translation. (**A**) Analogous compounds (**B**) Capped and polyadenylated Renilla mRNA (9 ng) was incubated with the supplemented TCH extract (2.6 μg/μL) in the presence of 40 μM of the compounds shown in (**A**) for 2 hours in a final volume of 8 μL, followed by luminescence detection. Each point represents a mean of triplicate measurements. The horizontal bar indicates the mean, and the vertical bar the SEM with p < 0.0001 (red), p < 0.001 (orange), p < 0.01 (green), and non-significant (black) as determined by two-way ANOVA with Dunnet multiple comparisons. (**C**) Inhibition of Renilla activity by the indicated compound concentrations as in (**A**). Each point represents the means and SEM of triplicate measurements in three independent experiments. Asterisks indicate significant inhibition (p < 0.05) as determined by a two-way ANOVA test with Tukey comparisons. **(D)** Luminescence assays made with 2 µL of parasite extract, 2 µL of mRNA (22 ng) and 1 µL of compounds to 40 µM. Significant decreases in luminescence were obtained by one-way ANOVA with Dunnet post-test relative to the DMSO sample with p = 0.0001 (***) and p < 0.0001 (****).

To determine if our assay could be further miniaturized, we conducted the assay with the best inhibitors in low-volume 384-well plates, in 5 µl. This later volume is chosen because it is the most often used volume in 1536-well ultra-high throughput screening assays. In this assay we verified a clear inhibition by compound 39 (Figure 5D).

### In vivo inhibition of protein synthesis by selected compounds

To conclusively demonstrate that our confirmed hit compounds truly inhibit protein synthesis in living parasites, we conducted a puromycylation assay, in which puromycin was incorporated into the elongating protein chains (Aviner, 2020; David et al., 2012). We treated exponentially proliferating epimastigotes with selected compounds for one hour prior to the addition of trace amounts of puromycin. As shown in Figure 6A, C39, C43, C39D, C39G, and C39H at 40 µM inhibited puromycin incorporation into elongating chains, indicating these compounds do indeed inhibit protein synthesis in live wild-type parasites. A quantitative analysis conducted in three distinct experiments indicates considerable suppression by these agents (Figure 6B). The results demonstrate that our in vitro translation assay is capable of identifying compounds that block *T. cruzi* protein synthesis in living parasites.

**Figure 6.**
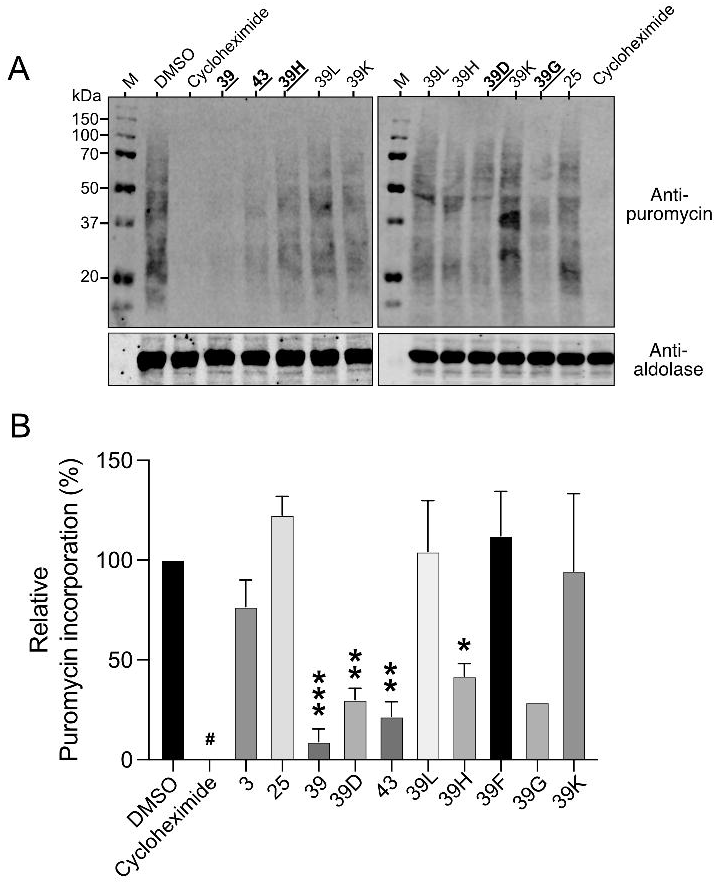
Effect of selected compounds on the puromycylation in intact *T. cruzi* epimastigotes. (**A**) Epimastigotes at 1 x 10^7^/mL were incubated in LIT medium containing 10% FBS for 1 hour at 28°C with the indicated compounds at 40 μM. Puromycin was added to each sample to 10 μg/mL, and after 15 minutes at 28°C, the parasites were collected by centrifugation, washed once with PBS, and lysed with RIPA buffer to estimate the amount of protein per assay. The samples were then mixed with SDS-PAGE sample buffer and used for Western blots probed with mouse anti-puromycin and rabbit anti-aldolase as loading controls and detected using anti-IgG IRDye 680 and 800, respectively. Markers (M) were added to the left side of each gel, and the respective size was indicated in kDa. (**B**) Quantitative analysis of puromycin incorporation relative to the incorporation in the sample containing only DMSO in the region between 15 and 35 kDa, from 3 to 4 independent experiments, except for compound 39G, tested in only one experiment. The values are means and SEM, with p < 0.001 (***), p < 0.005 (**), and p < 0.05 (*) estimated by one-way ANOVA with Dunnet test relative to the sample with DMSO.

## Discussion

We established an ultra-high-throughput *in vitro* translation assay to identify novel *T. cruzi* protein synthesis inhibitors. We utilized extracts from a genetically modified parasite exhibiting decreased hemin incorporation relative to wild-type strains to overcome the critical barrier to the establishment of this assay. The ability to reinitiate translation using *in vitro* transcribed RNA in the lysates of the mutant strain stems from a significant reduction in hemin following cell lysis; however, the precise mechanism underlying the inhibition of translation by hemin remains unknown.

We also confirmed that efficient in vitro translation requires capping at the 5’ end and polyadenylation at the 3’ end as well as the presence of a spliced leader because the mRNA without the spliced leader or additional nucleotides between the spliced leader and 7mGTPppp mRNA cap could not be translated. The cap added by the vaccinia enzyme is cap1, as opposed to the cap0 of most mammalian mRNA (where 7m is the only methylation) or the Cap4 of trypanosomes (Mair et al., 2000; Perry et al., 1987). Currently we do not know whether parasite extracts contain active methylating enzymes and thus further modify the reporter mRNA or if Cap1 is sufficient for translation by *T. cruzi*.

In this paper we optimized the incubation duration, RNA concentrations, DMSO concentration, and extract concentrations to obtain an assay with high performance. Furthermore, we miniaturized the assay to minimize the reagent quantities, which should facilitate screening of very large compound libraries. Furthermore, the translation was completely obliterated by general eukaryotic protein synthesis inhibitors. Interestingly, paromomycin, which preferentially inhibits *Leishmania* protein synthesis, had no effect on *T. cruzi* protein synthesis in vitro or in vivo. Despite the similarities between *Leishmania* and *T. cruzi*, as both belong to the Trypanosomatidae family, we were surprised that paromomycin did not inhibit *T. cruzi* translation. Furthermore, we noticed that paromomycin did not impact *T. cruzi* proliferation (data not presented). These data strongly support our contention that *T. cruzi*-specific protein synthesis inhibitors can be identified and developed.

In these proof-of-concept studies, we utilized agents previously shown to inhibit T. cruzi proliferation and compounds from DNDi, envisioning a high probability of finding structures that inhibit protein synthesis given the prevalence of protein synthesis inhibitors among anti-microbial, anti-mycotic, anti-malarial, and anti-leishmanial agents. Furthermore, these compounds were already shown to show very low risk of having toxicity to mammalian cells (Ahyong et al., 2016; Dmitriev et al., 2020; Hilal-Dandan & Brunton, 2016; O’Sullivan et al., 2018). After a series of follow-up tests, we determine the initial hit rate to be ∼6% with a confirmed hit rate of ∼50%. The initial hit rate is significantly higher than the high-throughput assays (0.3-1%), indicating significant enrichment. We found that two similar quinazoline compounds (C39 and C97) were highly effective as inhibitors of *T. cruzi* protein synthesis. In the follow-up studies, freshly purchased (in powder form) and prepared C39 was significantly more active than it was in the initial test.

Quinazolines, including C39, inhibit p21-activated kinase 4 (PAK4), which is a serine/threonine kinase involved in various cellular processes (Ramos-Alvarez et al., 2020). However, we have not found any publication connecting PAK4 to translation machinery. Similar compounds were found to target multiple stages of *Plasmodium falciparum,* the malaria-causing parasite, identified in high-throughput screenings (Gilson et al., 2017). Some quinazolines target dihydrofolate reductase (Guan et al., 2005), but it has not been shown that this enzyme has any impact on translation. In addition, some studies indicated that certain quinazoline derivatives exhibit activity against *T. cruzi*. A 14 quinazoline-2,4,6-triamine derivative and compounds with nitrobenzoyl substituents at 6-position demonstrated significant anti-trypanosomal activity against both epimastigote and trypomastigote forms while exhibiting low toxicity to human cells (Vazquez et al., 2024). Other studies also found that quinazolinone and quinazoline derivatives might be used against *T. cruzi* (Bollini et al., 2019). Therefore, our study may suggest that these compounds are inhibiting protein synthesis. It would be necessary to investigate whether quinazoline acts directly to inhibit components of translation machinery or indirectly through some proteins that play a role in translation initiation, elongation, or termination both in intact cells and cell extracts.

The width and breadth of commercially available chemical structures we tested do not allow for the reasoned structure-activity relationship analysis at this point. Synthesis and testing of focused quinazoline libraries is needed for this purpose.

## Conclusion

In summary, we developed a highly efficient ultra-high-throughput assay for screening *T. cruzi* protein synthesis inhibitors for the development of novel therapeutics for Chagas’ disease. Our studies suggest that certain quinazoline derivatives may be able to inhibit *T. cruzi* protein synthesis, but much work remains.

## Acknowledgments

We would like to thank Ayla Beatriz de Oliveira Santos, who helped in setting up the puromycylation assay, and Jadel Muller Kratz for providing DNDi compounds. This work was supported by an Exploratory/Developmental Research Grant (5R21AI154196) from the National Institutes of Health (NIH) to B.H.A. and partially by the Thematic Research Project from the Fundação de Amparo à Pesquisa do Estado de São Paulo (FAPESP), Process (2020/7870-4). S.S. received a fellowship from the Conselho Nacional de Desenvolvimento Científico e Tecnológico (CNPq, 303788/2020-8). C.C.M. was a recipient of a doctoral fellowship from the Coordenação de Aperfeiçoamento de Pessoal de Nível Superior (CAPES), Process (8887.920638/2023-0). G.R.R.S received a fellowship from the Conselho Nacional de Pesquisas (CNPq), Process 249194/2013-9.

