## Supplementary materials for "Ultra-high throughput *in vitro* translation assay for identification of *Trypanosoma cruzi-specific* protein synthesis inhibitors"

#### Obtaining TCH mutants

TCH mutants were obtained by a transfection of *T. cruzi* DM28c parasite strain previously transfected with pLEW-T7Cas9-Neo plasmid (Pacheco-Lugo et al., 2017), and positively selected with geneticin G418. To prepare the sgRNA DNA templates two oligonucleotides corresponding to the 5' end (SG5VPS23) and to the 3' end of the Vps23 gene (SG3VPS23) were used in a PCR reaction with the common primer (Supplementary Table S2). The donor was amplified using oligonucleotides P1VPS23 and P7VPS23, which anneal to the pPOT-mNeoGreen-Hygromycin B (Costa et al., 2018) and with the 5 ' and 3' regions of the VPS23 gene. Thereafter the cells were diluted in medium and maintained in the presence of 250 µg/mL of hygromycin B and 250 µg/mL of geneticin G418 (Invivogen, USA). One clone growing in hygromycin B did not contain the hygromycin B gene, which was not detected by PCR amplification similar to the parental line (T7Cas9), using the primers HygroF and HygroR and the total template DNA extracted from the cells, while two hygromycin B resistant clones presented the expected band of 550 base pairs (Figure S1A). Furthermore, the hygromycin was not inserted in the Vps23 locus compared to clones 8 and 9 as seen by a PCR using an hygromycin B primer and a primer to the 3' region of the Vps23 (Figure S1B). These observations were confirmed by sequencing an Illumina genomic library of TCH compared to the parental line. No changes in the relative number of hits of the Vps23 gene was observed in the TCH and only three hits to hygromycin sequences, compared to more than 100 of Vps23 were found.

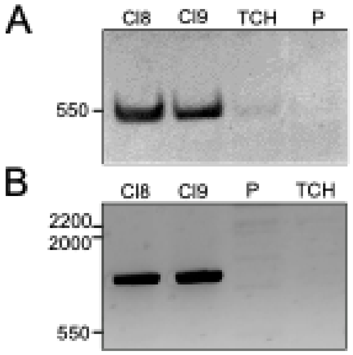

**Figure S1. PCR analysis of TCH mutant.** Total DNA was extracted from three clones growing in LIT medium containing 250 µg/mL hygromycin B. Each DNA sample was subjected to PCR (30 s at 94°C, 30 s at 57°C, and 30 s at 72°C). using in **(A)**, oligos HygroF and HygroR and HugroF and VPSUTRRev2 in **(B).**

### Table S1 – Compounds used in this study

| ompound Numb | lecular Weig | Inhibition % | PubChem CID | Catalog Number | Vendor/Source | MolPort | SMILES | Commercial name |
|---|---|---|---|---|---|---|---|---|
| 1 | 338.53 | -6.30 | 2973742 | 7957119 | ChemBridge Corporation | Molport-001-573-266 | C(CN1CCN(CCOC2CCCCC2)CC1)OC1CCCCC1 | |
| 2 | 419.92 | -6.93 | 2935107 | 7329145 | ChemBridge Corporation | Molport-002-121-505 | OC(=O)C(O)=O.CC1CN(CCOCCSc2ccc(Cl)cc2)CC(C)O1 | |
| 3 | 519.91 | 22.72 | 2834414 | 5217489 | ChemBridge Corporation | Molport-002-136-871 | [Cl-].CC(C)C1CCC(C)CC1OC(=O)Cn1c(-c2ccc(Br)cc2)[n+](C)c2ccccc12 | |
| 4 | 405.54 | -10.41 | 2920952 | 6965682 | ChemBridge Corporation | Molport-001-969-941 | CCCCC(=O)NC(c1ccc(cc1)N(CC)CC)c1ccc2cccnc2c1O | |
| 5 | 361.48 | -11.87 | 1378292 | 5431371 | ChemBridge Corporation | Molport-002-150-196 | COc1cc(CN(C)CCc2ccccc2)ccc1OCc1ccccc1 | |
| 6 | 383.56 | -7.28 | 16191881 | 21634851 | ChemBridge Corporation | Molport-005-030-842 | CCn1nc(C)c(CN2CCCC(C2)C(=O)c2cccc(OC(C)C)c2)c1C | |
| 7 | 372.48 | -3.36 | 24791021 | 57053236 | ChemBridge Corporation | Molport-005-103-493 | CN(CC1CCCN(CCc2ccc(F)cc2)C1)C(=O)c1cc(C)oc1C | |
| 8 | 310.48 | 2.78 | 2846021 | 5428477 | ChemBridge Corporation | Molport-002-149-870 | CC(=C)C1CCC(=CC1)CN2CCN(CC2)CC3=CC=CC=C3 | |
| 9 | 415.55 | -15.60 | 2954907 | 7790405 | ChemBridge Corporation | Molport-002-269-176 | CC(NS(=O)(=O)c1ccc(CCC(=O)N2CCN(C)CC2)cc1)c1ccccc1 | |
| 10 | 345.93 | -1.56 | 2992992 | 9008791 | ChemBridge Corporation | Molport-002-303-782 | Cl.CC1CN(CCOCCSc2ccc(C)cc2)CC(C)O1 | |
| 11 | 422.58 | -15.92 | 23723545 | 39103820 | ChemBridge Corporation | Molport-005-070-432 | CCN(CC1CCCN(CCc2cccc(F)c2)C1)Cc1ccc(cc1)C#CCCO | |
| 12 | 432.56 | 5.05 | 24790624 | 44284733 | ChemBridge Corporation | Molport-005-080-487 | CC(C)Cc1ccc(CN2CCCC(C2)NC(=O)c2nn(C)c(=O)c3ccccc23)cc1 | |
| 13 | 396.50 | -13.45 | 16188744 | 60081780 | ChemBridge Corporation | Molport-005-108-504 | CC(=O)c1cccc(CN2CCCC(CCC(=O)NCc3ccccc3F)C2)c1 | |
| 14 | 428.52 | 3.43 | 16191730 | 72292575 | ChemBridge Corporation | Molport-005-127-706 | COc1ccc(\C=C\CN2CCCC(CCC(=O)NCc3ccc(F)c(F)c3)C2)cc1 | |
| 15 | 355.48 | -8.17 | 16192471 | 88357737 | ChemBridge Corporation | Molport-005-150-468 | CCCn1ncc(CN2CCCC(C2)C(=O)c2cccc(OC)c2)c1C | |
| 16 | 410.53 | -1.11 | 16188753 | 83335963 | ChemBridge Corporation | Molport-005-143-669 | COc1ccccc1\C=C\CN1CCCC(CCC(=O)NCc2ccccc2F)C1 | |
| 17 | 396.53 | -7.25 | 23724036 | 44013371 | ChemBridge Corporation | Molport-005-079-988 | CCc1ccc(CN2CCCC(CNC(=O)c3cccc(OC)c3OC)C2)cc1 | |
| 18 | 326.91 | -0.46 | 2835280 | 5228728 | ChemBridge Corporation | Molport-002-111-924 | Cl.CN(C)c1ccc(CNCCC2CCOC(C)(C)C2)cc1 | |
| 19 | 375.55 | -12.00 | 16192086 | 59806838 | ChemBridge Corporation | Molport-005-108-055 | Cc1ccsc1C(=O)C1CCCN(Cc2cnc(s2)N2CCCC2)C1 | |
| 20 | 323.43 | -11.57 | 2838654 | 5269778 | ChemBridge Corporation | Molport-002-112-771 | CCOC(=O)C1CCCN(Cc2ccc(cc2)-c2ccccc2)C1 | |
| 21 | 307.86 | 2.32 | 9549450 | STK069023 | Vitas-M Laboratory, Ltd. | Molport-009-766-255 | Cl.CN(C)CC#CCC(O)(C1CCCC1)c1ccccc1 | |
| 22 | 393.42 | 5.61 | 2872175 | STK838714 | Vitas-M Laboratory, Ltd. | Molport-000-717-887 | CCOC(=O)c1c(NC(=O)Cn2cnc(n2)[N+]([O-])=O)sc2CC(C)CCc12 | |
| 23 | 273.72 | -7.56 | 5369911 | STK446818 | Vitas-M Laboratory, Ltd. | Molport-001-032-805 | C\C(=N/NC(=O)c1ccc(Cl)cc1)c1ccccn1 | |
| 24 | 339.32 | 9.03 | 6603435 | STK047082 | Vitas-M Laboratory, Ltd. | Molport-009-766-785 | Br.CCCCCCn1c2CCCc2c(=N)c2CCCc12 | |
| 25 | 395.42 | 68.77 | 16192607 | STK042510 | Vitas-M Laboratory, Ltd. | Molport-009-766-853 | Br.CCCCCCCCCn1c2CCCc2c(=N)c2CCCCc12 | |
| 26 | 367.40 | 7.63 | 2945620 | STK238653 | Vitas-M Laboratory, Ltd. | Molport-000-752-604 | CN(C)CCNc1ccc(cc1[N+]([O-])=O)-c1nn(C)c(=O)c2ccccc12 | |
| 27 | 379.39 | 4.48 | 1570351 | STK346036 | Vitas-M Laboratory, Ltd. | Molport-000-712-899 | CCOC(=O)c1c(NC(=O)Cn2cnc(n2)[N+]([O-])=O)sc2CCCCc12 | |
| 28 | 338.32 | -3.87 | 5405081 | STK101931 | Vitas-M Laboratory, Ltd. | Molport-000-804-060 | [O-][N+](=O)c1ccc(\C=N\NC(=O)CNc2ccc3ccccc3c2)o1 | |
| 29 | 324.76 | -1.43 | 5402622 | STK215250 | Vitas-M Laboratory, Ltd. | Molport-000-419-616 | Clc1ccccc1-c1ccc(\C=N\NC(=O)c2ccccc2)o1 | |
| 30 | 338.32 | 14.65 | 5393811 | STK107462 | Vitas-M Laboratory, Ltd. | Molport-000-787-111 | [O-][N+](=O)c1ccc(\C=N\NC(=O)CNc2cccc3ccccc23)o1 | |
| 31 | 359.45 | 1.47 | 1867212 | STK285784 | Vitas-M Laboratory, Ltd. | Molport-000-864-210 | CC(=O)N(Cc1cccnc1)c1cccc(c1)-c1nc2ccccc2s1 | |
| 32 | 321.53 | 2.69 | 5757429 | STK860027 | Vitas-M Laboratory, Ltd. | MolPort-001-936-684 | C\C(CC1CC2=C(CC1(C)C)C(C)(C)CCC2)=N/NC(N)=S \|c:5\| | |
| 33 | 334.64 | -0.58 | 2942060 | STK182767 | Vitas-M Laboratory, Ltd. | MolPort-002-091-101 | Cl.CN1CCC(CC1)OC(=O)c1ccc(Br)cc1 | |
| 34 | 313.46 | -3.18 | 1998473 | STK588424 | Vitas-M Laboratory, Ltd. | MolPort-002-607-403 | CN(C)CCCSc1nc(nc2CCCc12)-c1ccccc1 | |
| 35 | 414.59 | 6.60 | 16190843 | 32982674 | ChemBridge Corporation | Molport-019-805-226 | CC(C)n1nc(C)c(CN2CCCC(CCC(=O)NCc3ccccc3F)C2)c1C | |
| 36 | 398.50 | -0.46 | 16192204 | 96073626 | ChemBridge Corporation | MolPort-005-160-799 | COC(=O)c1ccc(CN2CCCC(CCC(=O)NCc3ccc(C)o3)C2)cc1 | |
| 37 | 422.56 | 7.85 | 16188908 | 89923774 | ChemBridge Corporation | Molport-005-152-546 | COc1cccc(CNC(=O)CCC2CCCN(C\C=C\c3ccccc3OC)C2)c1 | |
| 38 | 394.43 | 5.92 | 24817139 | 53804118 | ChemBridge Corporation | Molport-005-097-887 | CN(CC1CCCN(CCc2cccc(c2)C(F)(F)F)C1)C(=O)c1ccco1 | |
| 39 | 386.88 | 23.12 | 2951909 | 7749944 | ChemBridge Corporation | Molport-002-266-033 | Cl.COc1cccc(Nc2nc(NCC3CCCO3)c3ccccc3n2)c1 | |
| 40 | 429.99 | 4.79 | 23723620 | 13433324 | ChemBridge Corporation | MolPort-005-007-490 | COc1ccc(CN2CCCC(C2)N2CCN(CC2)c2cccc(Cl)c2)cc1OC | |
| 41 | 383.49 | -7.75 | 989046 | STL328189 | Vitas-M Laboratory, Ltd. | MolPort-000-657-600 | CN(C)CCNc1c(Cc2ccccc2)c(C)c(C#N)c2nc3ccccc3n12 | |
| 42 | 379.71 | -10.29 | 12005685 | STL041955 | Vitas-M Laboratory, Ltd. | MolPort-000-762-024 | [Cl-].Clc1ccc(Cn2cc(-c3ccccc3)[n+]3CCCc23)cc1Cl | |
| 43 | 410.32 | 40.10 | 24761168 | STL058959 | Vitas-M Laboratory, Ltd. | MolPort-000-727-433 | [I-].CN(C)c1ccc(cc1)-c1sc2cc(C)ccc2[n+]1C | |
| 44 | 379.39 | -4.10 | 3574126 | STL329752 | Vitas-M Laboratory, Ltd. | MolPort-001-506-876 | COC(=O)c1c(NC(=O)Cn2cnc(n2)[N+]([O-])=O)sc2CC(C)CCc12 | |
| 45 | 414.47 | 4.83 | 137054709 | STL495723 | Vitas-M Laboratory, Ltd. | MolPort-044-426-462 | C1CN(CCO1)c1nc(N\N=C\c2c[nH]c3ccccc23)nc(Nc2ccccc2)n1 | |
| 46 | 423.55 | -13.43 | 24816911 | 57359687 | ChemBridge Corporation | MolPort-005-104-032 | COc1ccccc1CN1CC(CCC1=O)C(=O)NCCN(C)CCc1ccccc1 | |
| 47 | 342.22 | 2.38 | 43234 | STL563469 | Vitas-M Laboratory, Ltd. | Molport-001-812-568 | CCCC1COC(Cn2cncn2)(O1)c1ccc(Cl)cc1Cl | |
| 48 | 347.54 | 34.82 | 3432274 | STL349484 | Vitas-M Laboratory, Ltd. | Molport-002-638-527 | CC(C)Oc1ccc(CNCCC2(CCOC(C)(C)C2)C(C)C)cc1 | |
| 49 | 403.75 | -4.17 | 16682470 | STL057843 | Vitas-M Laboratory, Ltd. | Molport-000-756-562 | [Br-].Cc1ccc(cc1)-c1cn(Cc2cccc(Cl)c2)c2CCC[n+]12 | |
| 50 | 361.55 | -7.82 | 132761820 | STL526173 | Vitas-M Laboratory, Ltd. | MolPort-002-514-195 | C[C@]12CCC3C(CC=C4CC(O)CC[C@]34C)C1CC\C2=N\NC(N)=S \|r,t:7\| | |
| 51 | 310.87 | -6.10 | 16192902 | STL512705 | Vitas-M Laboratory, Ltd. | Molport-009-757-382 | Cl.CN(C)CCCCCN(C)Cc1cc2ccccc2o1 | |
| 52 | 459.59 | 1.61 | 658950 | STK525017 | Vitas-M Laboratory, Ltd. | Molport-002-554-814 | CN1CCc2c(C1)sc1c2c(=O)n(-c2ccccc2)c2nnc(SCc3ccccc3)n12 | |
| 53 | 331.27 | -1.28 | 1247737 | STK464173 | Vitas-M Laboratory, Ltd. | MolPort-001-614-742 | CCc1sc(cc1Br)C(=O)NC1CCN(C)CC1 | |
| 54 | 358.35 | -3.55 | 5579166 | STK041579 | Vitas-M Laboratory, Ltd. | Molport-001-520-378 | Cc1ccc(NC(=O)CCC(=O)N\N=C\c2ccc(o2)[N+]([O-])=O)cc1C | |
| 55 | 372.38 | -6.80 | 4160908 | STK316177 | Vitas-M Laboratory, Ltd. | MolPort-002-795-985 | Cc1cc(NC(=O)C23CC4CC(C2)CC(C4)(C3)n2cnc(n2)[N+]([O-])=O)no1 | |
| 56 | 410.90 | -4.01 | 24747354 | STK194693 | Vitas-M Laboratory, Ltd. | MolPort-009-759-589 | OC(=O)C(O)=O.Clc1ccc(CN2CCCC(C2)C(=O)N2CCCCC2)cc1 | |
| 57 | 367.40 | -5.68 | 4462716 | STK316706 | Vitas-M Laboratory, Ltd. | MolPort-002-796-430 | [O-][N+](=O)c1ncn(n1)C12CC3CC(CC(C3)(C1)C(=O)Nc1ccccc1)C2 | |
| 58 | 421.48 | 0.41 | 23723846 | 12654542 | ChemBridge Corporation | Molport-005-004-981 | Fc1ccccc1N1CCN(CC1)C1CCCN(Cc2cccc(c2)C(F)(F)F)C1 | |
| 59 | 405.52 | -2.89 | 23723488 | 19837306 | ChemBridge Corporation | MolPort-005-026-035 | CSCCC(=O)N(C)CC1CCCN(CCc2cccc(c2)C(F)(F)F)C1 | |
| 60 | 434.98 | 5.00 | 23723925 | 26082236 | ChemBridge Corporation | MolPort-005-041-877 | COc1cc(CN(C)CC2CCCN(CCc3ccccc3F)C2)cc(Cl)c1OC | |
| 61 | 481.77 | -12.18 | 2922233 | 7001994 | ChemBridge Corporation | MolPort-002-232-003 | OC(=O)C(O)=O.CN1CCN(CCOCCOc2c(Cl)cc(C)cc2Br)CC1 | |
| 62 | 400.49 | 3.35 | 16190618 | 74383023 | ChemBridge Corporation | MolPort-005-130-810 | CCOc1cc(CN2CCCC(CNC(=O)c3ccc(F)cc3)C2)ccc1OC | |
| 63 | 347.21 | -6.44 | 4135073 | STK774949 | Vitas-M Laboratory, Ltd. | MolPort-002-725-547 | OC(COc1ccc2cc(Br)ccc2c1)Cn1ccnc1 | |
| 64 | 299.36 | -1.03 | 1990828 | BAS 01947743 | ASINEX Corporation | | CCOc1ccc(CNCCc2ccc3OCOc3c2)cc1 | |
| 65 | 358.35 | 4.50 | 4278357 | AK-968/40732169 | Specs | Molport-002-796-483 | [O-][N+](=O)c1ncn(n1)C12CC3CC(CC(C3)(C1)C(=O)Nc1ccon1)C2 | |
| 66 | 323.30 | 0.28 | 772084 | 5344177 | ChemBridge Corporation | Molport-001-507-210 | [O-][N+](=O)c1ccc(o1)C(=O)NN(c1ccccc1)c1ccccc1 | |
| 67 | 414.51 | 0.52 | 7208637 | E677-0419 | ChemDiv Inc. | Molport-007-752-919 | CN(C)CCCN(C(=O)c1nc2ccccc2s1)c1nc2ccc(F)cc2s1 | |

| | | | | | | | | |
|---|---|---|---|---|---|---|---|---|
| 68 | 454.60 | 5.74 | 2520 | V4629 | Sigma-Aldrich | Molport-000-721-258 | CC(C)C(CCCN(C)CCC1=CC(=C(C=C1)OC)OC)(C#N)C2=CC(=C(C=C2)OC)OC | |
| 69 | 274.40 | 8.14 | 1617333 | BAS 02788887 | ASINEX Corporation | Molport-001-983-473 | CN(C)CCCCCN(C)Cc1cc2ccccc2o1 | |
| 70 | 152.11 | 11.08 | 135403646 | HY-B1468 | MedChemExpress LLC | Molport-009-199-364 | Nc1nc2nn[nH]c2c(=O)[nH]1 | |
| 71 | 269.44 | 10.57 | 5475033 | 3182-0185 | ChemDiv Inc. | Molport-001-933-728 | CCN1CCS\C1=C/C=C/C1=[N+](CC)CCS1 | |
| 72 | 853.90 | 1.69 | 36314 | S1150 | Selleck Chemicals | Molport-001-742-627 | CC1=C2[C@H](C(=O)[C@@]3([C@H](C[C@@H]4[C@]([C@H]3[C@@H]([C@@](C2(C)C)(C[C@@H]1OC(=O)[C@@H]([C@H](C5=CC=CC=C5)NC(=O)C6=CC=CC=C6)O)O)OC(=O)C7=CC=CC=C7)(CO4)OC(=O)C)O)C)OC(=O)C | |
| 73 | 260.41 | -1.42 | 3613648 | 5422897 | ChemBridge Corporation | Molport-045-902-601 | CC(CCC=C(C)C)CN(C)CCc1ccccn1 | |
| 74 | 347.20 | 0.37 | 1502421 | STK774949 | Vitas-M Laboratory, Ltd. | | O[C@@H](COc1ccc2cc(Br)ccc2c1)Cn1ccnc1 | |
| 75 | 249.30 | 2.75 | 932871 | ST5024410 | TimTec LLC | Molport-002-006-131 | COc1ccc(cc1)C(=O)OC1CCN(C)CC1 | |
| 76 | 399.18 | 6.93 | 5579159 | STK040003 | Vitas-M Laboratory, Ltd. | Molport-001-520-373 | [O-][N+](=O)c1ccc(\C=N\NC(=O)CCC(=O)Nc2ccc(Cl)cc2Cl)o1 | |
| 77 | 623.69 | 47.54 | 6669960 | | Harvard | | CC(=O)n1cc([C@@H]2C[C@H](OCc3ccc(CO)cc3)OC(=C2)C(=O)N2CCN(Cc3ccc4OCOc4c3)CC2)c2ccccc12 | |
| 78 | 383.52 | 8.95 | 16190867 | | Broad Institute | Molport-005-157-774 | CCCn1ncc(CN2CCCC(C2)C(=O)c2cccc(OC(C)C)c2)c1C | |
| 79 | 443.41 | 14.04 | 1747861 | AK-968/15605994 | Specs | Molport-001-677-014 | [O-][N+](=O)c1cc(cc(c1)[N+]([O-])=O)C(=O)NCc1cn(nc1-c1ccccc1)-c1ccccc1 | |
| 80 | 330.38 | 18.93 | 886392 | OSSL_102421 | PrincetonBio | Molport-002-578-945 | Cc1ccc(Nc2nc(NCc3ccco3)c3ccccc3n2)cc1 | |
| 81 | 327.69 | -3.31 | 5002100 | C163-0471 | ChemDiv Inc. | Molport-001-604-259 | [O-][N+](=O)c1ncn(CCCC(=O)Nc2ccc(F)c(Cl)c2)n1 | |
| 82 | 442.90 | -11.55 | 4719406 | 4292 | Tocris Bioscience | Molport-000-861-927 | COc1ccc(CCNCc2ccc(OCc3ccc(Cl)nc3)c(OC)c2)cc1OC | |
| 83 | 369.50 | -10.49 | 16191661 | 99888991 | ChemBridge Corporation | Molport-005-165-579 | CCn1ncc(CN2CCCC(C2)C(=O)c2cccc(OC(C)C)c2)c1C | |
| 84 | 293.46 | -9.14 | 11836237 | AG-664/42673465 | Specs | Molport-002-806-907 | CC1CC(C)CN(CCOCCSc2ccccc2)C1 | |
| 85 | 440.53 | -11.50 | 3238629 | F0688-0091 | Life Chemicals | Molport-003-050-157 | C1CN(CCN1C2=CC=C(C=C2)F)C(CNS(=O)(=O)C3=CC=CC=C3)C4=CN=CC=C4 | |
| 86 | 372.49 | -6.54 | 16188996 | 23711140 | ChemBridge Corporation | Molport-005-036-204 | CSc1ccc(CN2CCCC(CNC(=O)c3ccc(F)cc3)C2)cc1 | |
| 87 | 376.47 | -8.90 | 1206996 | ST5010581 | TimTec LLC | Molport-001-967-329 | Cc1ccc(cc1)C(=O)c1c(NC(=O)c2ccncc2)sc2CCCCc12 | |
| 88 | 283.41 | 14.05 | 16954 | STL058959 | Vitas-M Laboratory, Ltd. | Molport-002-507-697 | CN(C)c1ccc(cc1)-c1sc2cc(C)ccc2[n+]1C | |
| 89 | 369.50 | -6.73 | 16187238 | 98254896 | ChemBridge Corporation | Molport-005-163-527 | CC(C)Oc1cccc(c1)C(=O)C1CCCN(Cc2cnn(c2)C(C)C)C1 | |
| 90 | 410.60 | -7.61 | 7507743 | F2018-0878 | Life Chemicals | | CCc1ccc2nc(sc2c1)N(CCN(C)C)C(=O)c1ccc2ncsc2c1 | |
| 91 | 445.46 | -10.37 | 2856632 | 5627014 | ChemBridge Corporation | Molport-000-467-957 | CC1(C)CC(C)(c2ccccc2)c2ccccc2N1C(=O)c1cc(cc(c1)[N+]([O-])=O)[N+]([O-])=O | |
| 92 | 272.25 | -12.50 | 3558505 | STK057185 | Vitas-M Laboratory, Ltd. | Molport-001-548-973 | [O-][N+](=O)c1ccc(o1)C(=O)N1CCc2ccccc2C1 | |
| 93 | 268.33 | -6.20 | 1547813 | F0013-0044 | Life Chemicals | Molport-000-519-032 | CC(=O)Nc1nc(cs1)-c1cccc2ccccc12 | |
| 94 | 364.38 | -22.91 | 2872283 | BAS 924218 | ASINEX Corporation | Molport-000-702-454 | CC1CCc2c(C1)sc(NC(=O)Cn1cnc(n1)[N+]([O-])=O)c2C(N)=O | |
| 95 | 548.67 | -1.50 | 16187141 | | Harvard | | OCc1ccc(CO[C@H]2C[C@H](C=C(O2)C(=O)N2CCN(Cc3ccc4OCOc4c3)CC2)C2CCCCC2)cc1 | |
| 96 | 299.38 | 5.86 | 6620905 | 6253-0534 | ChemDiv Inc. | Molport-007-570-276 | CN1C=C\C(=C/c2cc[n+](C)c3ccccc23)c2ccccc12 | |
| 97 | 320.38 | 57.49 | 3151039 | BAS 4913698 | ASINEX Corporation | Molport-001-998-854 | C(Nc1nc(Nc2ccccc2)nc2ccccc12)C1CCCO1 | |
| 98 | 414.48 | -11.93 | 16191862 | 84660490 | ChemBridge Corporation | Molport-005-145-469 | CC(=O)c1cccc(CN2CCCC(CCC(=O)NCc3ccc(F)c(F)c3)C2)c1 | |
| 99 | 333.43 | -11.84 | 648300 | 6892449 | ChemBridge Corporation | Molport-000-437-645 | CN(C)CCNc1c2CCCCc2c(C#N)c2nc3ccccc3n12 | |
| 100 | 432.51 | -3.30 | 3237664 | E155-0239 | ChemDiv Inc. | Molport-003-050-006 | COc1ccc(cc1)N1CCN(CC1)C(CNC(=O)Oc1ccccc1)c1cccnc1 | |
| 101 | 617.17 | 16.98 | 24892024 | F2019-1901 | Life Chemicals | Molport-009-688-040 | COc1cc2nc(sc2cc1OC)N(CCCN(C)C)C(=O)c1ccc(cc1)S(=O)(=O)N1CCc2ccccc12 | |
| 102 | 500.62 | 2.06 | 16187179 | | Harvard | | OCCCCO[C@H]1C[C@H](C=C(O1)C(=O)N1CCN(Cc2ccc3OCOc3c2)CC1)C1CCCCC1 | |
| 103 | 287.29 | 0.20 | SCYX0001613515 | | DNDi | | CC1CS(=O)(=O)CCN1\N=C\c2oc(cc2)[N+](=O)[O-] | Nifurtimox |
| 104 | 407.57 | -7.18 | DNDI0002233249 | | DNDi | | CCCCCCCCCCCCCCCCOP(=O)([O-])OCC[N+](C)(C)C | Miltefosine |
| 105 | 700.78 | -4.65 | SCYX0001433768 | | DNDi | | CC[C@@H]([C@H](C)O)N1N=CN(C1=O)c2ccc(cc2)N3CCN(CC3)c4ccc(OC[C@@H]5CO[C@](Cn6cncn6)(C5)c7ccc(F)cc7F)cc4 | Posaconazole |
| 106 | 473.40 | -5.03 | SCYX0001459208 | | DNDi | | CC(C)\N=C\1/C=C2N(c3ccc(Cl)cc3)c4ccccc4N=C2C=C1Nc5ccc(Cl)cc5 | Clofazimine |
| 107 | 705.63 | -0.08 | SCYX0001443738 | | DNDi | | CCC(C)N1N=CN(C1=O)c2ccc(cc2)N3CCN(CC3)c4ccc(OCC5COC(Cn6cncn6)(O5)c7ccc(Cl)cc7Cl)cc4 | Itraconazole |
| 108 | 926.06 | 93.90 | DNDI0002266587 | | DNDi | | C[C@@H]1OC(=O)C[C@H](O)C[C@H](O)CC[C@@H](O)[C@H](O)C[C@H](O)C[C@]2(O)C[C@H](O)[C@H]([C@H](C[C@@H](O[C@@H]3O[C@H](O)[C@@H](O)[C@H](N)[C@@H]3O)\C=C\C=C\C=C\C=C\C=C\C=C\C=C\[C@H](C)[C@@H](O)[C@H]1C)O2)C(=O)O | Amphotericin B |
| 109 | 343.51 | -4.48 | SCYX0001713235 | | DNDi | | CCN(CC)CCCCCCNc1cc(OC)cc2c(C)ccnc12 | Sitamaquine |
| 110 | 340.43 | -7.74 | DNDI0003296706 | | DNDi | | NC(=N)c1ccc(OCCCCCOc2ccc(cc2)C(=N)N)cc1 | Pentamidine |
| 111 | 389.25 | 21.41 | DNDI0003579092 | | DNDi | | | |
| 112 | 416.49 | -2.43 | DNDI0003286690 | | DNDi | | | |
| 113 | 429.41 | -7.50 | DNDI0003295996 | | DNDi | | | |
| 114 | 339.05 | -1.42 | SCYX0001186759 | | DNDi | | | |
| 115 | 367.38 | -1.31 | DNDI0002491044 | | DNDi | | Fc1ccc(O[C@@H]2CCN(C2)C(=O)Nc3cc(n[nH]3)c4ccccn4)cc1 | DNDI-1044 |
| 116 | 335.13 | 1.81 | SCYX0002166148 | | DNDi | | Cc1c(nnn1c2ccccn2)C(=O)Nc3ccc4COB(O)c4c3 | DNDI-6148 |
| 117 | 279.32 | -2.39 | SCYX0000749397 | | DNDi | | CSc1ccc(OCc2ncc([N+](=O)[O-])n2C)cc1 | Fexinidazole |
| 118 | 367.10 | -9.69 | SCYX0001267158 | | DNDi | | | |
| 119 | 447.53 | -3.44 | DNDI0003578526 | | DNDi | | | |
| 120 | 370.33 | -5.42 | DNDI0002490690 | | DNDi | | [O-][N+](=O)c1cn2CC[C@H](COc3ccc(nc3)c4ccc(F)cc4)Oc2n1 | DNDI-0690 |
| 121 | 445.44 | 9.00 | DNDI0003095561 | | DNDi | | FC(F)(F)c1cnn(C[C@@H]2CCN(C2)C(=O)Nc3[nH]nc(c4ccccn4)c3C5CC5)c1 | DNDI-5561 |
| 122 | 307.31 | 7.58 | DNDI0003366174 | | DNDi | | Cc1nc(N)nc2[nH]c(C)c(c3ccc(C#N)c4OCOc34)c12 | DNDI-6174 |
| 123 | 381.63 | 13.55 | SCYX0001608210 | | DNDi | | | |
| 124 | 359.26 | 22.78 | SCYX0001730347 | | DNDi | | | |
| 125 | 296.30 | 22.65 | DNDI0003296000 | | DNDi | | | |

| | | | | | | | | |
|---|---|---|---|---|---|---|---|---|
| 126 | 368.31 | 18.98 | DNDI0003296749 | | DNDi | | | |
| 127 | 293.32 | 20.46 | DNDI0003286588 | | DNDi | | | |
| 128 | 260.25 | 20.42 | SCYX0001268526 | | DNDi | | [O-][N+](=O)c1nccn1CC(=O)NCc2ccccc2 | Benznidazole |
| 25E | 297.24 | 50.30 | 16682138 | STK054125 | Vitas-M Laboratory, Ltd. | Molport-009-765-813 | Br.CCn1c2CCCc2c(=N)c2CCCCc12 | |
| 25F | 465.56 | 71.70 | 52997752 | STK039469 | Vitas-M Laboratory, Ltd. | Molport-009-766-746 | Br.CCCCCCCCCCCCCCn1c2CCCc2c(=N)c2CCCCc12 | |
| 39A | 374.84 | 65.40 | 2952932 | 7769487 | ChemBridge Corporation | Molport-002-267-092 | Cl.Fc1cccc(Nc2nc(NCC3CCCO3)c3ccccc3n2)c1 | |
| 39B | 372.85 | 57.60 | 2953078 | 7772685 | ChemBridge Corporation | Molport-002-267-254 | Cl.Oc1ccc(Nc2nc(NCC3CCCO3)c3ccccc3n2)cc1 | |
| 39C (97C) | 321.38 | 53.50 | 50945469 | 9222426 | ChemBridge Corporation | Molport-016-588-602 | C(Nc1nc(Nc2ccncc2)nc2ccccc12)C1CCCO1 | |
| 39D | 374.84 | 71.90 | 2927619 | 7125070 | ChemBridge Corporation | Molport-002-238-369 | Cl.Fc1ccc(Nc2nc(NCC3CCCO3)c3ccccc3n2)cc1 | |
| 39F (97F) | 308.81 | 45.90 | 52995972 | STL162970 | Vitas-M Laboratory, Ltd. | Molport-009-762-133 | Cl.CN(C)c1nc(NCC2CCCO2)c2ccccc2n1 | |
| 39G (97B) | 350.42 | 59.10 | 2929218 | 7174729 | ChemBridge Corporation | Molport-001-998-856 | COc1ccccc1Nc1nc(NCC2CCCO2)c2ccccc2n1 | |
| 39H (97H) | 338.38 | 54.50 | 2929638 | 7184299 | ChemBridge Corporation | Molport-002-241-534 | Fc1ccccc1Nc1nc(NCC2CCCO2)c2ccccc2n1 | |
| 39I(97I) | 324.81 | 38.20 | 52995693 | STL162695 | Vitas-M Laboratory, Ltd. | Molport-009-761-261 | Cl.OCCNc1nc(NCC2CCCO2)c2ccccc2n1 | |
| 39J(97J) | 229.28 | 34.90 | 22330544 | STL491425 | Vitas-M Laboratory, Ltd. | Molport-008-958-620 | C(Nc1ncnc2ccccc12)C1CCCO1 | |
| 39K (97K) | 377.27 | 55.10 | 45928522 | STK843001 | Vitas-M Laboratory, Ltd. | Molport-009-767-252 | Cl.Clc1cccc(Nc2nc(N3CCOCC3)c3ccccc3n2)c1 | |
| 39L(97D) | 316.79 | 60.60 | 9549592 | STL152186 | Vitas-M Laboratory, Ltd. | Molport-003-356-377 | Cl.OCCNc1nc(Nc2ccccc2)nc2ccccc12 | |
| 39M (97E) | 371.87 | 46.90 | 52995827 | STL157846 | Vitas-M Laboratory, Ltd. | Molport-009-761-663 | Cl.C(Nc1nc(NCc2ccccn2)nc2ccccc12)C1CCCO1 | |
| 39E (129) | 334.78 | 50.90 | 135402554 | STK975722 | Vitas-M Laboratory, Ltd. | Molport-002-101-106 | Cc1cc(=O)[nH]c(SCc2nc(no2)-c2ccc(Cl)cc2)n1 | |

Table S2 – Primers used in this project

| Oligonucleotide | Sequence |
| --- | --- |
| SG5VPS23 | 5´GAAATTAATACGACTCACTATAGGGGGACACGGGACCAGCT GTGGTTTTAGAGCTAGAAATAGC |
| SG3VPS23 | 5’GAAATTAATACGACTCACTATAGGTTTTTTTTTTTCTATGTGTA GTTTTAGAGCTAGAAATAGC |
| Common | 5'AAAAAAGCACCGACTCGGTGCCACTTTTTCAAGTTGATAACG GACTAGC. |
| P1VPS23 | 5'GGATTTTTTATTATTAATATTTTTTTTTAAGGTATAATGCAGACCT GCTGC |
| P7VPS23 | 5’TATGTATATATTCACACAGAGACAGACCCACCGGAACCACTAC CAGGAACC |
| VPSUTRRev2 | 5’GCGAGTTTCCCTCCTTATCCG |
| HygroF | 5’GAAAAAGCCTGAACTCACCGC |
| HygroR | 5’GCAAACTGTGATGGACGACAC |
